# Isohydric species show earlier drought-induced declines in stem water content, rehydration, sap flow, and growth than anisohydric species

**DOI:** 10.1101/2025.07.30.666311

**Authors:** Sharath S. Paligi, Christina A. Hackmann, Jan Schick, Michela Audisio, Heinz Coners, Martina Mund, Christian Ammer, Christoph Leuschner

## Abstract

- Isohydric species reduce water potential fluctuations through more stringent stomatal regulation. It is unclear whether isohydric behavior leads to smaller reversible diurnal stem shrinkage (i.e. greater nocturnal stem rehydration) during drought and a lower drought sensitivity of radial growth compared to anisohydric behavior.
- With synchronous high-resolution sap flow and dendrometer measurements in mature anisohydric European beech and isohydric Douglas fir trees in pure and mixed stands, we quantified declines in stem water content (SWC), stem rehydration, sap flow and radial growth during soil dry-down and determined the critical soil moisture levels (expressed as Relative Extractable Water, REW) and elapsed desiccation time until 30-90% reductions in these traits.
- Sap flow and growth started to decline in both species at REW ∼0.6, preceding declines in stem rehydration. Water-spending Douglas fir approached 50% drops in SWC, sap flow, stem rehydration and growth during soil dry-down faster than beech, indicating higher drought sensitivity. In mixture, both species reached these reduction levels later than in monoculture, suggesting positive mixing effects on the species’ drought resistance.
- Our findings demonstrate that SWC, sap flow and radial growth decrease earlier than nocturnal stem rehydration, with isohydric and anisohydric species exhibiting different timelines of physiological downregulation during soil dry-down.

## Introduction

‘Global-change-type droughts’ are increasingly impacting forests worldwide, causing growth reduction, vitality loss and elevated tree mortality (Allen et al. 2015, Adams et al. 2017, McDowell et al. 2020, Breshears et al. 2021, Hammond et al. 2022). Drought-induced tree mortality is primarily driven by hydraulic failure, often exacerbated by pest outbreaks that exploit weakened trees (Choat et al. 2018, Arend et al. 2021, Hajek et al. 2022, McDowell et al. 2022). Forests in Central Europe have been particularly affected, with major timber species such as Norway spruce, Scots pine, oak species, and European beech (hereafter referred to as “beech”) experiencing substantial growth declines (Neycken et al. 2022, Weigel et al. 2023, Leuschner et al. 2023, Enderle et al. 2024, Gribbe et al. 2024, Leifsson et al. 2024) and increased tree mortality during recent hot droughts (Schuldt et al. 2020, Senf et al. 2021, Thonfeld et al. 2022, Knutzen et al. 2025). The exceptional sequence of severe droughts during 2018-2020 and in 2022 in Central Europe demonstrates the threats trees will be exposed to in a warming and drying climate (van der Woude et al. 2023, Gharun et al. 2024).

With intensifying climate extremes, proactive adaptive forest management through the selection of more drought- and heat-resistant species is essential to sustain forest ecosystem functions (Bauhus et al. 2017, Messier et al. 2022, Decarsin et al. 2024). With its high growth rates and reputed drought resistance, Douglas fir (*Pseudotsuga menziesii* (Mirb.) Franco), native to the Pacific Northwest of North America, has sparked the interest of Central European foresters as a viable option (Eilmann and Rigling 2012, Brus et al. 2019, Thomas et al. 2022). To reduce climate risks, foresters in Central Europe are nowadays widely preferring mixed species forests, such as combining native beech and introduced Douglas fir (Glatthorn et al. 2021, Topanotti et al. 2024). Both species differ not only in their leaf habit, phenology, and wood anatomy, but also in their stomatal regulation strategy, with likely consequences for drought resistance and resilience (Leuschner 2020, Leuschner and Meinzer 2024). Moreover, drought response traits are not only species-specific but can also be modulated by interspecific interactions (Paligi et al. 2024). Tree species mixtures have been found to alleviate, exacerbate, or have no effect on drought stress, depending on species and context (Pardos et al. 2021, Haberstroh and Werner 2022, Hackmann et al. 2024). To understand the underlying mechanisms of species interactions, knowledge about the relevant, species-specific functional traits is necessary.

Isohydric species tend to stringently close their stomata to maintain more stable leaf water potential, which often results in reduced carbon assimilation, whereas anisohydric species maintain gas exchange longer during water deficits but may risk hydraulic failure (Tardieu and Simonneau 1998, Meinzer et al. 2017). While beech shows a typical anisohydric behavior with considerable diurnal and seasonal leaf water potential fluctuations, Douglas fir pursues a more isohydric strategy with more stringent down-regulation of stomatal conductance upon water deficits (Leuschner et al. 2022, Schumann et al. 2024). Although the isohydric–anisohydric framework is widely used to categorize stomatal responses to drought, recent studies have shown that isohydric species are not necessarily more carbon-limited or quicker to close stomata, highlighting the need for a more nuanced, trait- and context-dependent perspective (Martínez-Vilalta and Garcia-Forner 2017, Kannenberg et al. 2019, Volaire 2018, Paligi et al. 2024).

Understanding how plants adjust transpiration and growth in response to soil drying is crucial for predicting their performance during drought (Joshi et al. 2022). Stomatal conductance is influenced by various factors, notably decreasing soil moisture, increasing atmospheric vapor pressure deficit, and decreasing leaf water potential (Buckley 2019), with tree species differing considerably in their responsiveness to these factors (Oren et al. 1999, Anderegg et al. 2017, Flo et al. 2021). A common paradigm of stomatal action postulates that stomata respond to decreasing water availability in a manner that balances assimilation efficiency and hydraulic safety (Sperry et al. 2017, Anderegg et al. 2018), such that stomatal regulation primarily reduces the risk of embolism formation, while maximizing carbon gain due to enhanced water-use efficiency (Yang et al. 2016, Guerrieri et al. 2019). However, recent studies suggest that stomatal regulation may be optimized to maintain adequate stem turgor for radial growth, rather than to avoid cavitation (Peters et al. 2023, 2025a, Potkay and Feng 2023). Stomatal closure not only decreases carbon gain and growth but also reduces the capacity of leaves to cool through transpiration (Marchin et al. 2016).

Soil moisture plays a crucial role for the gas exchange and growth of plants, as transpiration and photosynthesis start to decrease when soil moisture drops below a critical level (θ_crit_) and the available energy is increasingly partitioned from evapotranspiration towards sensible heat flux. This shifts the system from energy limitation to water limitation (Seneviratne et al. 2010, Fu et al. 2021, Denissen et al. 2022). Various attempts have been made to define θ_crit_ for forest ecosystems, with soil moisture availability expressed either in absolute (soil volumetric water content, VWC; or soil matric potential, Ψ_Soil_) or relative units (relative extractable water, REW) (Baldocchi et al. 2005, Fu et al. 2021). For example, Granier et al. (1999) found a consistent down-regulation of transpiration in various temperate forests at about 0.4 REW. θ_crit_ has been found to vary with soil texture and tree species (Granier et al. 1999, Brinkmann et al. 2016, Wankmüller et al. 2024), but our understanding of tree species differences in the response of plant water status, water consumption and growth to drying soil under comparable climatic and edaphic conditions is still incomplete.

Water shortage limits both the carbon sources (i.e., photosynthesis) and carbon sinks of plants (i.e. meristematic activity) (Fatichi et al. 2014, Mund et al. 2020, Martínez-Sancho et al. 2022). Wood growth requires sufficient turgor pressure in the cambium, which is reduced under both low soil moisture and high VPD (Steppe et al. 2015, Peters et al. 2021, Niessner et al. 2024). The reversible diurnal stem diameter fluctuations are used to calculate tree water deficit (TWD) as an indicator of stem dehydration and tree drought stress (Zweifel et al. 2016). Temperate tree species facilitate stem rehydration by means of stomatal regulation (Peters et al. 2023), with species characterized by lower hydraulic resistance in the stem (vessel-bearing angiosperms) exhibiting more effective stem rehydration overnight than tracheid-bearing conifers (Peters et al. 2023, Hackmann et al. 2024). Previous work has shown that Douglas fir stands with their usually higher stand basal area and leaf area consume more water (Paligi et al. 2025) and exhibit less effective night-time stem rehydration (Hackmann et al. 2024) than beech. However, it is not clear if and how stem dehydration, sap flow and stem radial growth are synchronized during soil drying.

By operating high-resolution sap flow sensors and automated dendrometer systems synchronously during a soil drying event in a severe summer drought, we examined the response of stem water content, stem rehydration, sap flow and radial growth of an isohydric (Douglas fir) and an anisohydric (beech) tree species to soil dry-down, both in pure and in mixed stands. Based on these observations, we aimed to better understand how water status- and growth-related traits are differently influenced by soil moisture depletion in species with loose or stringent stomatal regulation. Such knowledge is crucial for attempts to predict the survival probability of trees in a drier and warmer climate (Granier 1999, Groover et al. 2025, Senf et al. 2025) and thus may aid adapting forest management strategies to future conditions (Novick et al. 2024). The following hypotheses guided our research: (1) The two tree species differ in stand-level soil moisture depletion patterns, with water-demanding Douglas fir exhausting soil water reserves faster than beech. (2) Stem water status and growth progressively deteriorate during soil desiccation in both species. We expect that isohydric Douglas fir, with stricter stomatal control, will exhibit decline in sap flow (transpiration) at relatively higher REW to maintain higher stem water content longer (i.e., longer elapsed desiccation time). In contrast, anisohydric beech, with its flexible stomatal regulation, will sustain transpiration down to lower REW, and thus will reach critical stem water content and sap flow reduction thresholds (70% or 90%) earlier than Douglas fir (i.e., shorter elapsed desiccation time). (3) Isohydric Douglas fir maintains favorable stem water status (higher stem water content and lower tree water deficit) longer than beech, resulting in a greater drought resistance of radial growth. (4) Due to species complementarity in functional traits, both species reach critical water status and growth performance thresholds later in mixture than in the respective monocultures.

## Materials and methods

### Study area

The study was conducted in the Pleistocene lowlands of northern Germany in the Lüß forest in the Lüneburg Heath region, Lower Saxony. The area is characterized by a cool-temperate suboceanic climate with a mean annual temperature of 9.2 °C and an average precipitation of 785 mm (1993-2022 period, gridded monthly raster data was obtained from the German Weather service (DWD), accessed via the R package `rdwd v1.8.0 (`Boessenkool 2023)). The sites are located at 165 m a.s.l. on level terrain without access to groundwater. The soils are acidic, nutrient-poor Spodo-Dystric Cambisols developed in sandy-loamy unconsolidated substrate (79% sand, 15% silt and 6% clay in the top 30 cm of the mineral soil; Foltran et al. 2023).

Three experimental plots of 50 m x 50 m size were established at a maximum distance to each other of 300 m in even-aged mature stands, one plot each in stands dominated by beech or Douglas fir (with the target species contributing 93.5% and 85.4% of the basal area, respectively), and one mixed beech–Douglas fir plot (44.2% and 46.3% of basal area, respectively). The plots are hereafter referred to as pure beech, pure Douglas fir and mixed beech–Douglas fir plots. In the mixed stand, beech is referred to as ‘mixed beech’ and Douglas fir as ‘mixed Douglas fir’. The tree age of the cohort-like stands was 70 years (pure Douglas fir), 82 years (mixed Douglas fir), and 85 years (pure beech, mixed beech). Basal area was highest in the pure Douglas fir plot (49.1 m^2^ ha^-1^), followed by the mixed beech–Douglas fir (36.7 m^2^ ha^-1^) and the pure beech plot (28 m^2^ ha^-1^; Table 1). Understory vegetation was absent under the beech and beech–Douglas fir canopies, while a sparse shrub layer existed in the pure Douglas fir plot.

**Table 1.**
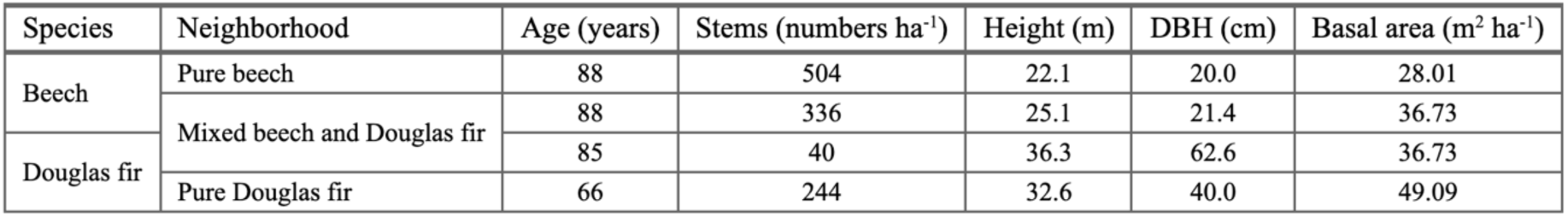
Stand structural characteristics of the three study plots.

### Soil moisture and microclimate monitoring

Volumetric soil water content (VWC) was monitored in the three plots at 5 cm, 20 cm, 50 cm and 100 cm depth with TDR sensors (CS650, Campbell Scientific, Shepshed, UK) at hourly intervals using a Campbell CR300 data logger. To compare the extent of soil moisture depletion during the growing season among the three plots, we expressed the variable soil moisture reserves in the profile to 130 cm depth as Relative Extractable Water (REW) (Granier et al. 1999), i.e. the fraction (1 – 0) of maximum soil water storage being available at a point in time. To derive REW, we first normalized volumetric water content (VWC) at each measurement depth (5, 20, 50, and 100 cm) using the minimum and maximum values observed between January 2019 and December 2022 (unpublished data). We then calculated available water (AW, in mm) for each soil horizon by multiplying these values by the corresponding layer thickness: 0–10 cm (5 cm depth), 11–30 cm (20 cm), 31–70 cm (50 cm), and 71–130 cm (100 cm). The four AW values were summed to estimate the total soil moisture reserve in the 0–130 cm profile, which was then expressed as a fraction of the maximum AW measured to obtain total profile REW.

Air temperature and relative air humidity were measured with a CS215 sensor, and precipitation with an ARG314 rain gauge (Campbell Scientific, Logan, UT, USA) at a weather station in an open area near the study plots. All variables were recorded using a CR300 datalogger. Vapor pressure deficit (VPD) was computed from temperature and air humidity using the magnus formula (Tetens 1930).

For quantifying tree-level drought responses, a tree’s response time to a stress event of known severity and duration can be measured, i.e. the time elapsed since the onset of stress until critical physiological thresholds are reached. The severity and duration of drought stress may be quantified as cumulated VPD over the period of desiccation. This can be expressed as ‘elapsed desiccation time’ (in VPD hours) by multiplying VPD (kPa) with the time (hours) elapsed to reach specific levels of REW (if mean VPD is 1 kPa, the unit VPD hours equals hours). This approach was previously applied in controlled greenhouse studies (e.g. Blackman et al. 2019, Paligi et al. 2024) and accounts for variability in environmental conditions during the measurement period, allowing to compare different treatments in their drought exposure more accurately.

We fit shape-constrained additive models (SCAM) to express the relationship between REW and elapsed desiccation time, with the aim to compare beech and Douglas fir in pure and mixed stands in their growth, water status and water consumption responses to soil desiccation (see Table 2 for model specifications and Supplementary information for detailed description). To compare the temporal progression of water stress in the two species and different neighborhoods (pure vs mixed) upon soil desiccation, we used the models to extract the desiccation time elapsed to reach REW levels of 0.9, 0.7, 0.5, 0.3 and 0.1 for each plot.

**Table 2.**
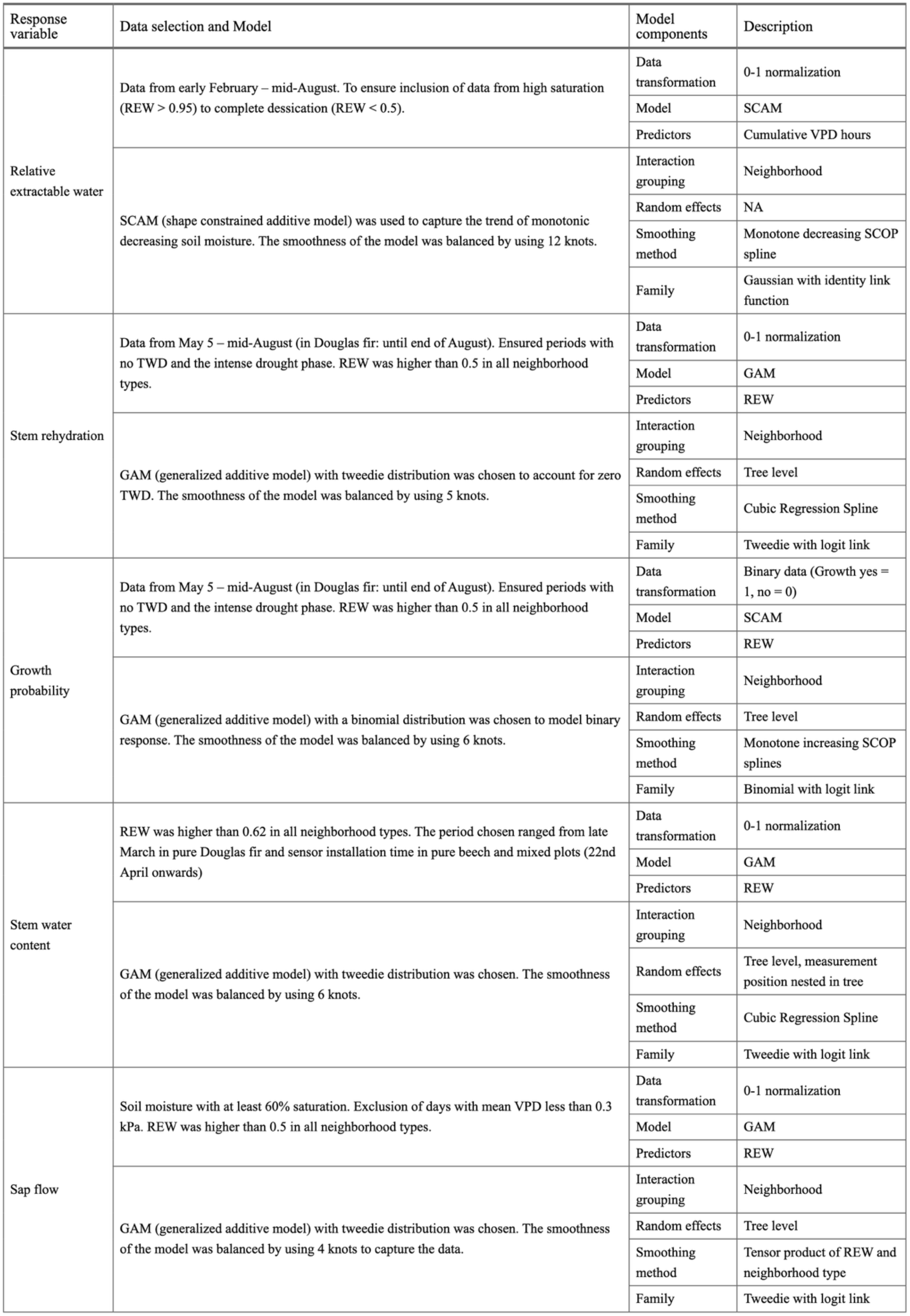
Summary of data selection and modeling approaches for analyzing the response of five physiological variables – relative extractable water, tree water deficit, growth probability, stem water content, and sap flow under different neighborhood conditions. The table describes the data filtering process, the selected model types and the key model components such as data transformation, predictor variables, random effects and smoothing parameters. The models account for tree species interactions. Missing values (NA) indicate instances where a specific model component was not applicable or not required.

### Measurement of radial stem growth and stem rehydration

Daily radial growth and daily stem rehydration were derived from stem diameter variation measurements. Stem diameter at breast height (1.3 m above ground) was measured at 10-min intervals with 16 band dendrometers (DC2 and DC3; Ecomatik, Munich, Germany) mounted on each four trees of the two species per stand. We used glide rings to minimize friction between wire and bark. The tree bark was smoothed where necessary, to ensure firm contact with the stem and reduce its influence on stem diameter variation measurements. The selected trees represented the typical diameter range of trees forming the upper canopy layer in the stands. Radial stem growth and tree water deficit (TWD, µm) were calculated according to the zero-growth concept (Zweifel et al. 2016) with the R package `treenetproc v0.1.4` (Haeni et al. 2020, Knüsel et al. 2021). The concept distinguishes between irreversible diameter growth (i.e., diameters exceeding the previous maximum diameter) and reversible swelling and shrinking of the stem due to changes in stem water content, where TWD quantifies how far the current diameter lies below the previous maximum diameter. On the diurnal scale, the minimal (predawn) TWD is of particular physiological interest, since it indicates a tree’s capacity to rehydrate the stem overnight and thus may reflect the potential to grow (Dietrich et al. 2018, Zweifel et al. 2021, Peters et al. 2023, 2025b, Hackmann et al. 2024). In accordance with this concept, we equate daily minimum TWD with the degree of “stem rehydration”, scaled from 0 to 1 for each tree. Thus, a stem rehydration of 1 signifies a daily minimum TWD of 0, i.e. full rehydration, while stem rehydration < 1 reflects a persisting TWD, i.e. incomplete stem rehydration overnight. Daily radial growth was treated as a binary variable, indicating whether or not a tree grew on a particular day.

### Measuring daily tree water use and stem water content

Sap flow sensor installation, data handling and sap flow upscaling procedures are described in detail in Paligi et al. (2025). In brief, in spring 2022, we installed 37 HPV-06 Dual-Method-type sap flow sensors (Implexx Sense, Melbourne, Australia) in 0.5 and 1.5 cm sap wood depth in the pure beech (n = 9) and pure Douglas fir (n = 9) plots, and in the mixed beech–Douglas fir plot (9 beech and 10 Douglas fir trees). To ensure correct sensor positioning, we removed bark thicker than 0.5 cm and installed the sensors following Forster et al. (2019). Probe alignment was checked initially, and, if necessary, further time-dependent probe misalignment corrections were applied to account for seasonal changes in stem water content (Larsen et al. 2020). Sap wood properties were determined by extracting wood samples with a wood corer (Haglöf, Långsele, Sweden).

For upscaling sap flow from point measurements to the tree level, we additionally employed Heat-Field-Deformation sensors (HFD100, ICT International Pty Ltd., Armidale, Australia) to characterize the radial flow profiles in the sapwood of the measured trees. We adopted the approach of Caylor and Dragoni (2009), integrating time-dependent continuous measurements and time-independent radial flow profile measurements (for methodological details see Paligi et al. (2025)). The HFD sensors measured at multiple depths (either at 0.5 cm-intervals when the needle length was 3.5 cm, or at 1 cm-intervals when the needle length was 8 cm). To model maximum sapwood depth, we employed Generalized Additive Models (GAMs) with gamma distribution, accounting for species and depth. From these data, we calculated daily sap flow (L day^-1^) for each tree as described in Paligi et al. (2025).

#### Stem water content

Stem water content (SWC) was simultaneously recorded by the HPV-06 sap flow sensors at two measuring depths (0.5 cm and 1.5 cm in the sapwood). For our analysis, we extracted daily maximum (predawn) stem water content which typically occurs between 2:00 and 6:00 a.m. This period of low sap flow was chosen to avoid potential errors associated with the heat pulse technique and thermal conduction during high sap flow. We then averaged SWC over the outer and inner measuring positions.

#### Midday twig water potential

We conducted weekly measurements of midday twig water potential as a measure of foliar water status between 8 June and 23 August 2022. Measurements were taken from sun-exposed terminal twigs with 3-5 leaves for beech and two entire fascicles for Douglas fir, sampling all twelve trees using a Scholander pressure chamber (PMS Instruments, Corvallis, Oregon, USA). Samples were collected by tree climbing between 10 a.m. and 2:00 p.m. Measurements were conducted within 5 to 30 minutes of cutting the branches. If measurements could not be taken within 5 minutes, samples were wrapped in aluminum foil and sealed in zip-lock bags to prevent desiccation until measurement.

### Modeling physiological thresholds

To detect REW thresholds at which sap flow, radial growth, SWC and stem rehydration start to decline, we employed a suite of statistical models tailored to each response variable within the framework of Generalized Additive Models (GAMs) using the `R` package `mgcv v1.8-39`. The models used appropriate distributions (i.e., Gaussian, Tweedie or binomial) to account for the nature of each dataset (see Table 2 for model specifications, Fig S2-5 for visual assessments of model fit, and Supplementary Information for detailed model fitting procedures).

The models were fitted using relative data (normalized to 0 – 1) at the tree level, except for radial growth probability, which was treated as a binary response (0 or 1). The data spanned variable environmental conditions across the three stand types. We extracted REW and desiccation time thresholds from the fitted models which corresponded to 30%, 50%, 70% and 90% reductions in the physiological response.

Model selection for each variable was based on identifying the best-fitting model that most effectively captured the data. This involved testing different combinations of distribution families, and spline functions with varying levels of complexity (i.e. the number of knots, basis functions, etc.) to determine the optimal fit. The final model choices were further validated through diagnostic plots to ensure their suitability.

### Data analysis and visualization

All data handling and analysis was performed in R software v4.1.3 (R core team 2023) in the framework of the package `tidyverse v2.0.0` (Wickham et al., 2019). Figures were produced using the package `ggplot2 v3.4.0` (Wickham, 2016) and cowplot v1.1.1 (Wilke et al. 2020) and tables using `flextable v0.8.5` (Gohel and Skintzos 2023).

## Results

### Weather conditions and soil moisture

The 2022 growing season was at our site exceptionally dry and hot, with the main drought period lasting from mid-April to mid-August with only 128 mm precipitation (Figure 1). The total annual precipitation during 2022 was 555 mm which was ∼30% lower than the long-term mean (785 mm). This period included 21 hot days (maximum temperature >30°C) and reached a peak temperature of 41.1 °C in late July (Fig. 1a), approaching the highest recorded temperature in Germany. Hourly VPD exceeded 3 kPa on 14 days, peaking at 6.5 kPa. Soil moisture declined steadily from mid-April, when relative extractable water (REW) was 0.6 –0.8 in the profile, to mid-August with REW <0.05 (grey lines in Fig 1b-e).

**Figure 1.**
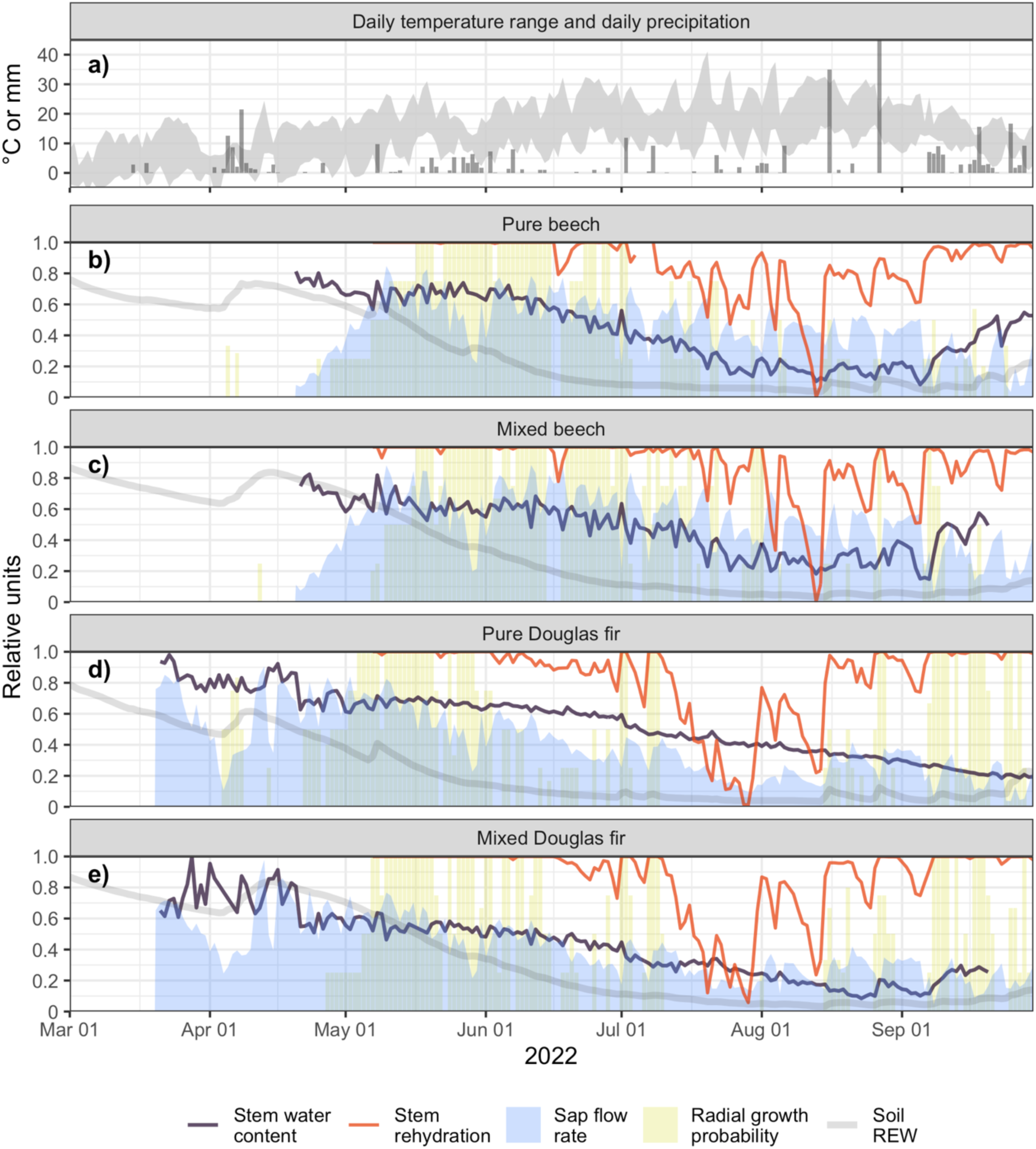
a) Daily temperature maxima and minima and daily temperature (shaded grey) and daily precipitation (grey bars) in the period early March – late September 2022. b) – e) Variation of daily sap flow (shaded blue), daily stem growth (yellow bars), daily minimum TWD (red line), daily maximum stem water content (SWC, black line) and daily mean REW (0-100 profile) for pure beech, mixed beech, mixed Douglas fir and pure Douglas fir in the period early March – late September 2022. All variables are given in relative terms (1 = highest value recorded, 0 = lowest value recorded). For radial growth, the probability of growth on that day according to a GAM is given.

### Water use, growth, and stem water status during progressive drought

During the extended drought from April to August 2022, daily sap flow, stem water content, and the frequency of days with radial growth decreased with advancing soil desiccation (Fig 1b-e) and decreasing foliar midday water potentials (Fig. S6) in both species and neighborhood constellations (pure and mixed). It was associated with a marked decrease in stem rehydration (orange lines in Figure 1b-e). At maximum drought in late July and August, radial growth ceased for several weeks in all stands (yellow bars in Fig. 1b-e). Mean tree-level water consumption was reduced in the pure beech stand to 18% of maximum recorded water consumption in this stand (blue area in Fig. 1b), and to even 8% in the pure Douglas fir stand (Fig. 1d). In the mixed beech-Douglas fir stand, beech reduced its water consumption during drought to 25% and Douglas fir to 8% of maximum water consumption in this stand (blue area in Fig. 1c and e). The midday water potential decreased (more negative) with decreasing soil REW, with Douglas fir in mixture showing the most negative and beech in mixture the least negative values (Figure S6).

Under ample water supply, beech and Douglas fir had similar mean stem water contents (SWC) in the pure and mixed stands, averaging around 35% in beech and 29% in Douglas fir (black lines in Fig. 1b-e). At peak drought, SWC decreased to 23% in pure beech and 28% in mixed beech, while in Douglas fir, it dropped to 20% in mixture and to 18% in the pure stand.

Daily stem rehydration was usually complete until early June in Douglas fir and late June in beech, but decreased greatly toward July and August (orange lines in Figure 1 b-e). During peak drought, daily minimum TWD reached 430 - 973 µm for Douglas fir, and 130 - 368 µm for beech. Accordingly, stem diameter growth was relatively low, cumulating to on average 3.1 and 1.9 mm for beech, and 3.2 and 3.1 mm for Douglas fir in pure and mixed stands, respectively, from May to mid-August. During this period (107 days), beech had on average 56 and 59 growth days, while Douglas fir had only 39 and 43 growth days in the pure and mixed stands, respectively. Notably, all trees showed a similar number of growth days (ca. 65) throughout the entire growing season, mainly driven by Douglas fir growth after drought release in late summer (yellow bars in Figure 1 b-e).

### Patterns of soil moisture depletion

During the spring/summer drought of 2022 (mid-April to mid-August), daily REW minima reached less than 0.04 in all three plots (Fig. 1b-e). Soil moisture depletion proceeded faster under the pure Douglas fir than under the pure beech and mixed beech-Douglas fir stands: It took 341 kPa hours to reduce REW from 0.9 to 0.5 in the profile under the Douglas fir stand, but 750 kPa hours and 772 kPa hours under the beech and the mixed stands, respectively (Fig. 2a and S1, Table 3). In contrast, more advanced soil water depletion (from REW 0.5 to 0.1) occurred faster under the pure beech stand (532 kPa hours) than under pure Douglas fir (725 kPa hours) and the mixed stand (744 kPa hours; Fig. 2b and S1, Table 3).

**Figure 2.**
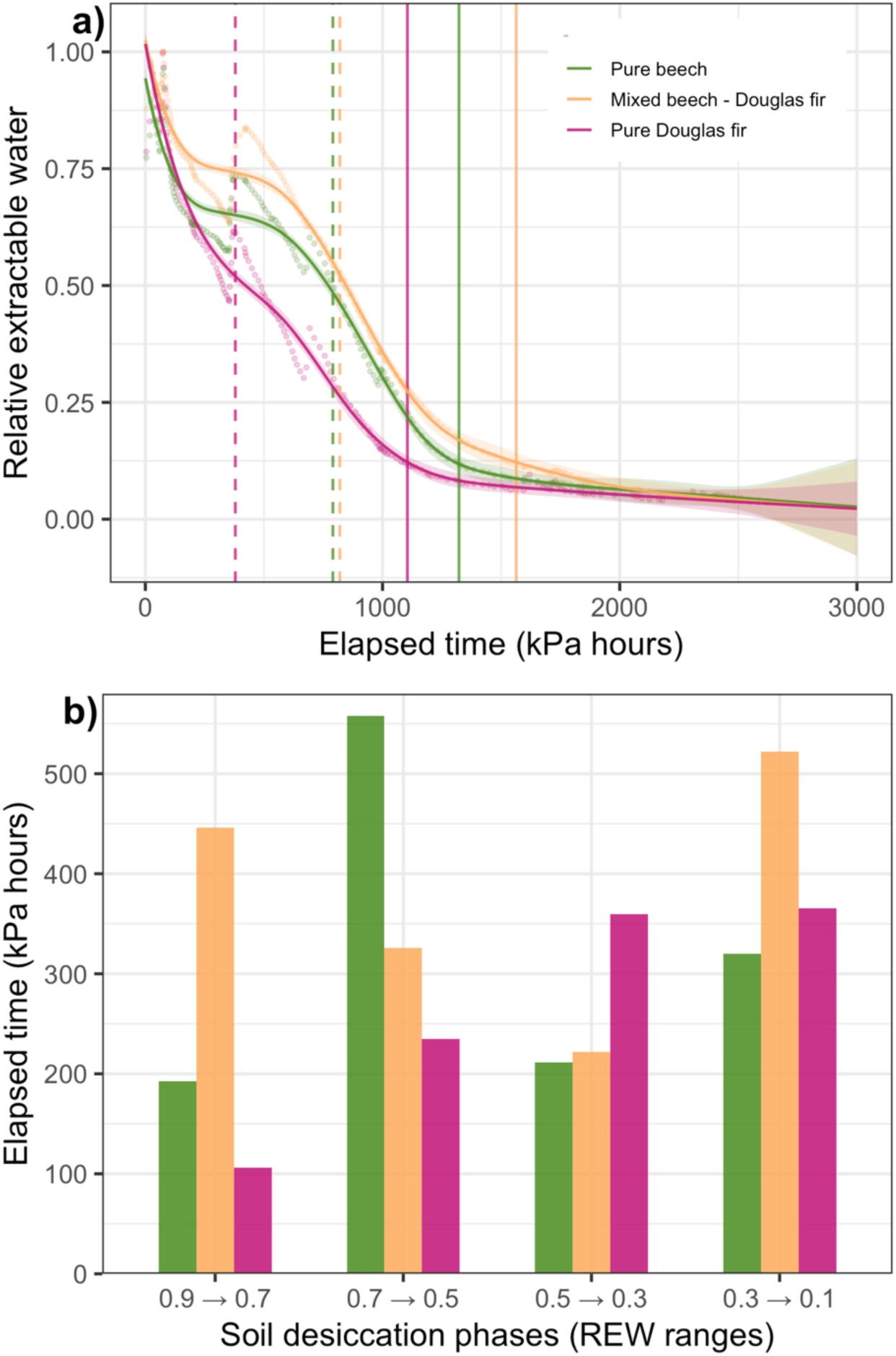
a) Soil moisture depletion curves during the drought in summer 2022 for the pure beech (green), pure Douglas fir (purple) and the mixed beech-Douglas fir stand (orange). Given is the decrease in relative extractable water (REW) as a function of elapsed desiccation time (expressed in kPa hours). The curves represent fitted Shape-Constrained Additive Models (SCAMs) with 95% confidence intervals (shaded regions) and data points (dots). Vertical lines denote the time spans needed to reach REW 0.5 (dashed lines) or REW 0.1 (solid lines) in the three stands. (b) Length of time passed during which soil moisture decreased from REW 0.9 to 0.7, from REW 0.7 to 0.5, from REW 0.5 to 0.3, or from REW 0.3 to 0.1 in the pure beech (green), pure Douglas fir (purple) and the mixed beech-Douglas fir stand (orange). The time span is expressed in kPa hours to normalize the length of the period of drought stress exposure by means of cumulated VPD.

**Table 3.** Time elapsed during the spring/summer drought 2022 (mid-April – late August) to reduce relative extractable soil water (REW) from 0.9 to 0.7, from 0.7 to 0.5, from 0.5 to 0.3, and from 0.3 to 0.1 in profiles under pure beech, pure Douglas fir and beech-Douglas fir mixture. The time interval is expressed in kPa hours (time interval x corresponding VPD level) to relate soil moisture depletion to the cumulative evaporative demand in that period

| Stand type | Time elapsed (kPa hours) to reduce REW for specified interval |  |  |  |
| --- | --- | --- | --- | --- |
|  | 0.9 → 0.7 | 0.7 → 0.5 | 0.5 → 0.3 | 0.3 → 0.1 |
| Pure beech | 192.77 | 557.72 | 211.53 | 320.05 |
| Mixed beech - Douglas fir | 446.08 | 326.13 | 221.98 | 521.81 |
| Pure Douglas fir | 105.83 | 234.88 | 359.94 | 365.49 |

### Critical soil moisture levels of tree water status, sap flow and radial growth

In the course of soil moisture depletion during the 2022 drought, sap flow started to decline in both species already at REW levels of 0.8 – 0.6 (Fig. S2). Fifty percent sap flow reduction was reached in Douglas fir at REW 0.13, but only at REW 0.07 in beech, indicating more sensitive stomatal regulation in Douglas fir (Fig. 3 and Fig. S2). Correspondingly, a sap flow reduction by 70% was observed in pure and mixed Douglas fir at REW 0.08 and 0.1, respectively, while in the pure beech stand, it occurred at considerably lower REW (<0.01). It is noteworthy that the maximum sap flow reduction observed during the drought was 70% in beech (in the pure stand), but reached 90% in Douglas fir (pure stand).

**Figure 3.**
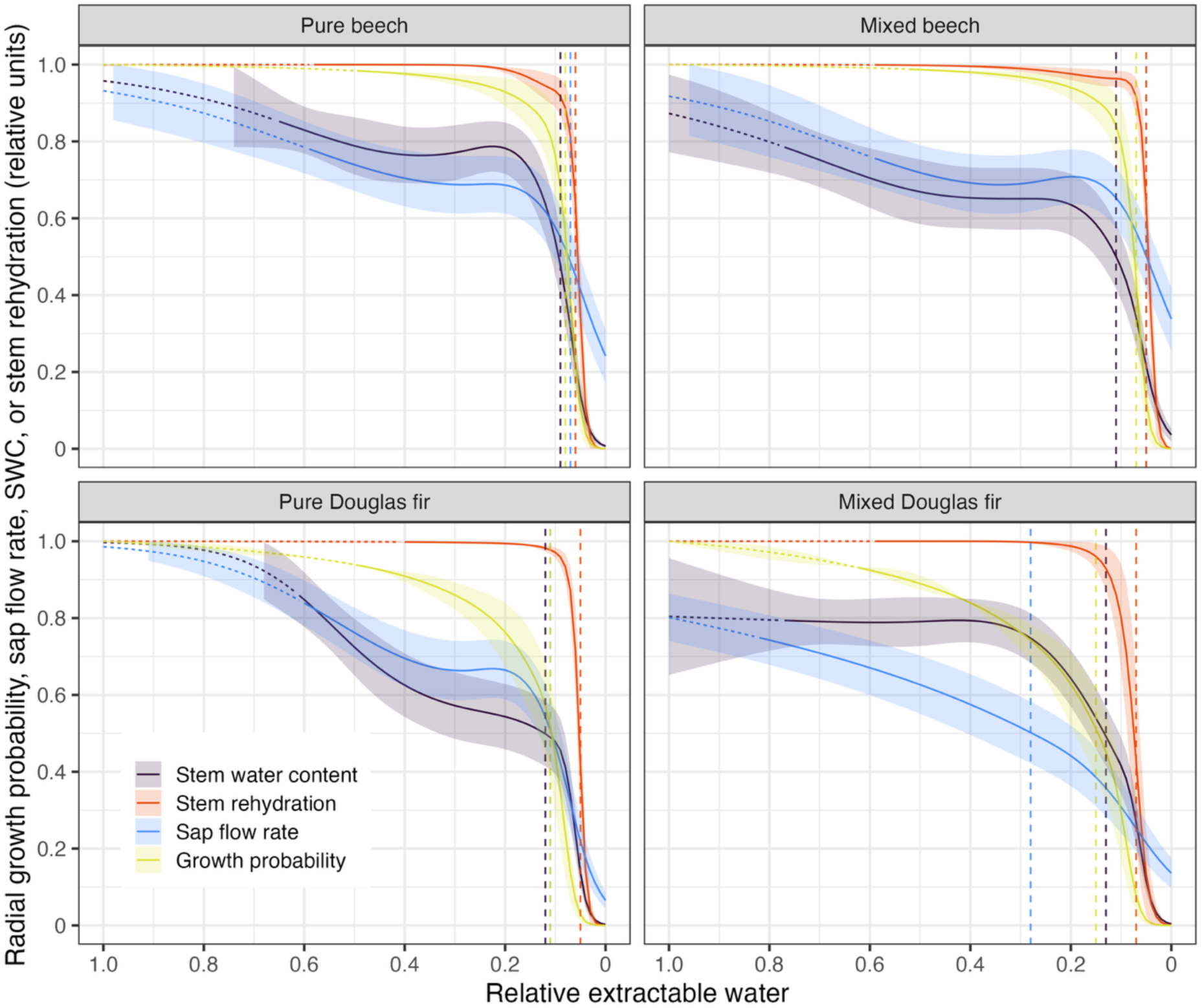
Daily sap flow (blue), daily mean stem water content (SWC, black), daily minimum tree water deficit (or stem rehydration, red), and probability of radial growth (green) in relation to REW for beech (top row) and Douglas fir (bottom row) in pure (left column) and mixed (right column) stands in the summer 2022. All variables are given in relative terms (1 = highest value recorded, 0 = lowest value recorded). For radial growth, the probability of growth on that day according to a GAM is given. The smoothed curves represent model fits (see Table 2 for models and model fits used for the different variables), with shaded regions indicating 95% confidence intervals. Vertical dashed lines denote the REW values at which 50% of the maximum values is reached for the four variables. Dotted segments of the curves indicate extrapolated model results beyond the observed REW range for that variable.

The sap flow decrease was closely followed by a reduction in SWC, which reached 50% of the maximum recorded at about REW 0.1 in beech and REW 0.13 in Douglas fir (Fig. 3 and Fig. S2). In Douglas fir, also radial growth (expressed as growth probability) began to decrease in the early phase of soil desiccation at REW 0.8 – 0.6, while growth impairment was recorded in beech only at REW <0.4. Accordingly, growth probability was reduced to 50% already at REW 0.14 and 0.2 in pure and mixed Douglas fir, while this happened in beech only at REW <0.1 (Fig. 3 and Fig. S3).

Stem rehydration was complete in beech and Douglas fir until REW fell below ∼0.2 (Fig. 3 and Fig. S4). Both species responded similarly to soil desiccation and reached 50% reduction in stem rehydration at ∼ REW 0.05. Thus, in the pure stands, 50% reductions in sap flow and growth were generally reached earlier in Douglas fir than in beech in the pure stands, but the species did not differ with respect to the REW thresholds for SWC and stem rehydration (Table 4 and Fig. S5 and S4). In the mixed stand, Douglas fir responded to soil desiccation more sensitively than beech with reductions in SWC, sap flow and growth, but not in stem rehydration. Moreover, Douglas fir appeared to be more sensitive in the mixed than the pure stand, while beech differed between the two stand types only little (Table 4).

**Table 4.** REW levels (as a fraction of 1) at which stem water content (SWC), stem rehydration, sap flow and radial growth of beech and Douglas fir in the pure and mixed stand were reduced during the 2022 drought by 30%, 50%, 70% or 90% of their recorded maxima in spring. In addition, the time elapsed to reach these reduction levels (‘desiccation time’, expressed in kPa hours by multiplying the time intervals with the corresponding VPD level) is given. NA – missing values or reduction level not observed. 3000+ - time interval exceeding 3000 kPa hours, but desiccation interrupted by rainfall.

| Variable | Neighborhood | REW at specified reduction levels |  |  |  | Desiccation time (kPa hours) |  |  |  |
| --- | --- | --- | --- | --- | --- | --- | --- | --- | --- |
|  |  | 30% | 50% | 70% | 90% | 30% | 50% | 70% | 90% |
| Stem water content | Pure beech | 0.18 | 0.10 | 0.07 | 0.04 | 1163 | 1436 | 1860 | 2652 |
|  | Mixed beech | 0.69 | 0.10 | 0.05 | NA | 592 | 1719 | 2304 | 3000+ |
|  | Mixed Douglas fir | 0.71 | 0.13 | 0.05 | NA | 543 | 1516 | 2304 | 3000+ |
|  | Pure Douglas fir | 0.50 | 0.13 | 0.07 | 0.02 | 422 | 1080 | 1538 | 2960 |
| Stem rehydration | Pure beech | 0.05 | 0.06 | 0.06 | 0.07 | 2386 | 2110 | 2110 | 1860 |
|  | Mixed beech | 0.04 | 0.05 | 0.05 | 0.06 | 2534 | 2304 | 2304 | 2123 |
|  | Mixed Douglas fir | 0.06 | 0.07 | 0.09 | 0.11 | 2123 | 1991 | 1799 | 1646 |
|  | Pure Douglas fir | 0.05 | 0.05 | 0.06 | 0.07 | 2082 | 2082 | 1789 | 1538 |
| Sap flow | Pure beech | 0.16 | 0.07 | 0.01 | NA | 1201 | 1860 | 3000+ | 3000+ |
|  | Mixed beech | 0.16 | 0.07 | 0.00 | NA | 1358 | 1991 | 3000+ | 3000+ |
|  | Mixed Douglas fir | 0.55 | 0.24 | 0.10 | NA | 790 | 1156 | 1719 | 3000+ |
|  | Pure Douglas fir | 0.25 | 0.13 | 0.08 | 0.00 | 838 | 1080 | 1364 | 3000+ |
| Growth probability | Pure beech | 0.13 | 0.10 | 0.07 | 0.03 | 1279 | 1436 | 1860 | 2891 |
|  | Mixed beech | 0.13 | 0.09 | 0.06 | 0.03 | 1516 | 1799 | 2123 | 2788 |
|  | Mixed Douglas fir | 0.25 | 0.20 | 0.15 | 0.09 | 1140 | 1234 | 1404 | 1799 |
|  | Pure Douglas fir | 0.17 | 0.14 | 0.11 | 0.07 | 978 | 1050 | 1152 | 1538 |

Despite similar REW levels at which given stem rehydration and SWC reductions were reached, the standardized time elapsed to approach 50%, 70% or 90% reductions in these variables differed markedly among the species and stands. Corresponding to the more sensitive downregulation of sap flow in Douglas fir, this species reached 50%, 70% or 90% reductions in sap flow much earlier than beech and also reduced growth more rapidly to these levels (Table 4). Species differences were weaker with respect to SWC and stem rehydration. In most cases, both species reached the thresholds later in the mixture than in the respective pure stands, suggesting a lower stress level.

The relation of stem rehydration with sap flow and SWC during the soil desiccation phase followed a sigmoidal shape. Sap flow of Douglas fir responded more sensitively than sap flow of beech, decreasing already at higher stem rehydration (Figure 4). In terms of SWC sensitivity to decreasing stem rehydration, the species differed much less. Interestingly, both species responded less sensitively in the mixed stand than in their respective pure stand (Figure 4).

**Figure 4.**
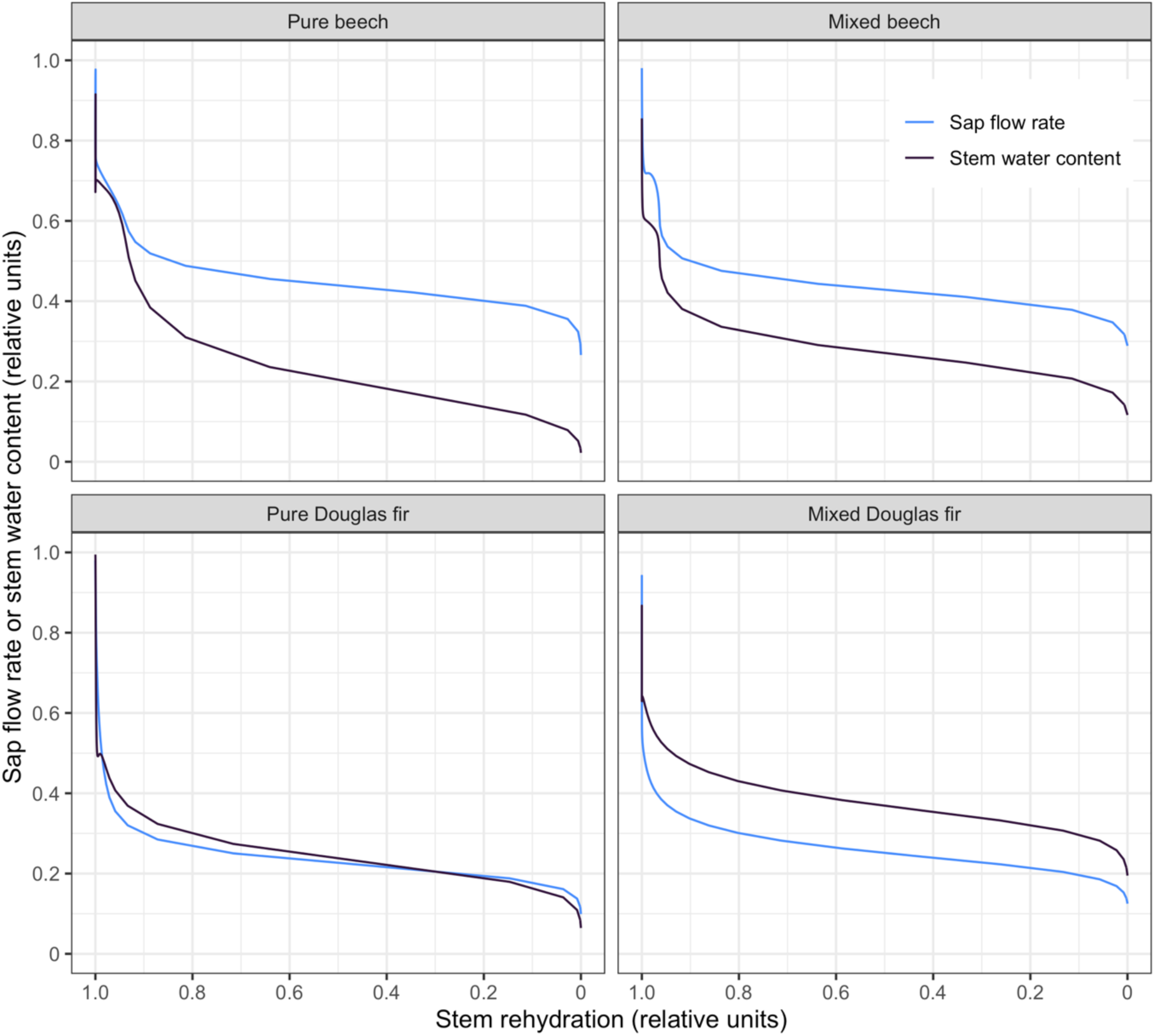
Dependence of daily sap flow (blue) and daily mean stem water content (SWC, black) on stem rehydration (or daily maximum tree water deficit, TWD) in beech (top row) and Douglas fir (bottom row) in pure (left column) and mixed (right column) stands in the summer 2022.

## Discussion

In this study, we present a comprehensive approach to investigate stem water status during soil drying. Our findings reveal relative extractable water (REW) thresholds corresponding to key physiological responses (i.e., 30, 50, 70 and 90% reductions in growth probability, sap flow, stem water content, and stem rehydration) across stand types. While Douglas fir and beech exhibited dissimilar REW thresholds corresponding to reductions in sap flow and growth, their responses in stem water content and stem rehydration were similar, with stem rehydration persisting to lower REW levels than did the other three variables. Notably, the elapsed desiccation time required to reach these thresholds varied considerably across stand types, emphasizing the role of species identity and stand properties in modulating drought resilience. By extending methodologies previously applied in greenhouse studies to natural forest settings, we quantify soil desiccation in terms of elapsed time, providing a more dynamic understanding of drought progression.

### Water use strategies and phenology determine soil moisture depletion patterns

As soil moisture declines, plant water status deteriorates, constraining plant metabolism and vitality and eventually leading to mortality (Granier et al. 1999, Choat et al. 2018, McDowell et al. 2022, Mendonca et al. 2023). Our study demonstrates that the velocity of soil moisture depletion differs between beech and Douglas fir stands under similar soil and climate conditions, and it is influenced by species mixing effects.

The pure Douglas fir stand reduced soil moisture more rapidly than did the pure beech stand, reaching 0.1 REW (i.e. 90% water depletion in the 0-100 cm profile) 215 kPa hours earlier, which confirms our first hypothesis. During the first phase of soil drying (REW decrease from 0.9 to 0.5), the velocity of soil moisture depletion under Douglas fir was markedly faster than under beech or in the mixed stand. This difference can be attributed to the higher water losses to the atmosphere during the growing season, driven by the higher basal area and large foliage area of Douglas fir (Leuschner and Meinzer 2024), as well as its earlier onset of transpiration in spring compared to deciduous beech (Thomas and Winner 2000, Moore et al. 2004, Link et al. 2014, Paligi et al 2025). In contrast to the first phase, the dry-down during the second phase (REW drop from 0.5 to 0.1) was faster under beech than under pure Douglas fir and the mixed stand. This relates to a generally lower stand-level water consumption of beech due to its smaller basal area (Paligi et al. 2025), but also to its sustained water extraction under further soil drying, reflecting a less pronounced reduction in stomatal conductance. The contrasting soil desiccation patterns reflect well the more isohydric water status regulation patterns of Douglas fir in comparison to the anisohydric behavior of beech (Meinzer et al. 2017, Schumann et al. 2024, Paligi et al. 2024, 2025, Leuschner et al. 2022, Leuschner and Meinzer 2024). This is consistent with ecosystem-level flux-tower observations that show a more stringent canopy conductance reduction upon drought in conifers compared to angiosperm trees (Gharun et al. 2020). Moreover, leaf minimum conductance after stomatal closure has been found to be ten times higher in beech than in Douglas fir (Paligi et al. 2024), which may explain the faster soil moisture depletion under beech later in the desiccation process. These patterns align with predictions of plant hydraulic models, which indicate that species-specific stomatal regulation stringency is a principal driver of soil desiccation dynamics (Preisler et al. 2023, Decarsin et al. 2024). The mixed stand took even longer than the pure beech stand to reach 0.1 REW, with desiccation patterns being more similar to the beech stand in the first phase and more resembling the Douglas fir stand in the second phase of soil drying.

### REW thresholds of critical stem water status

Despite a largely different wood anatomy, foliage phenology and hydraulic strategy, beech and Douglas fir showed critical relative reductions (50%, 70%, or 90%) in SWC and stem rehydration at comparable REW thresholds, and within a narrow REW range (0.13 - 0.03). Notably, SWC started to decline well ahead of stem rehydration, as demonstrated by the 30% reduction thresholds in Table 4 and Figure 3. While both variables target daily maximum (predawn) values of stem water content, and stem rehydration were measured with different sensors (at 0.5 and 1.5 cm depth in the sapwood vs. on the bark surface) and thus captured moisture changes in different stem tissues (only outer sapwood vs. total sapwood plus cambium, phloem and bark). It appears that the outer sapwood of beech and Douglas fir begins to lose moisture in the course of soil drying well before the capacity for night-time rehydration of other tissues is compromised. These results highlight the crucial role of stem water reserves for the buffering of drought impacts (Cermák et al. 2007, Köcher et al. 2013, Preisler et al. 2022) and suggest that different tissue types are prioritized differently during daily rehydration, potentially favoring cambial growth and phloem transport (Sevanto 2018, Salmon et al. 2019, Peters et al. 2023, 2025a).

In our stands, sap flow started to decline in this severe drought already at REW levels well above 0.6, much ahead of stem rehydration, and apparently in parallel with the decline in stem water content. Our sap flow data indicate for both beech and Douglas fir a down-regulation of transpirational water loss with soil drying in two steps: an initial slight reduction, which may begin already at REW ∼0.7, and a second sharp decline at REW ∼0.2, which coincides with the rapid decrease in stem rehydration. Our results differ from earlier reports about critical REW levels, where soil moisture begins to limit the maximum transpiration of temperate trees and the transpiration process shifts from energy-limited to moisture-limited (Seneviratne et al. 2010). For example, sap flow or energy balance measurements in oak, spruce and Douglas fir stands revealed REW levels of 0.3 – 0.4, at which soil drying started to reduce transpiration (Black 1979, Granier 1987, Biron 1994, Granier et al. 1999). We speculate that the earlier onset of sap flow reduction in our stands may be due to specific soil conditions. The lack of groundwater access and the sandy soil texture with its low water holding capacity and steep decrease in soil hydraulic conductivity during soil drying likely have triggered an earlier physiological response (Wankmüller et al. 2024). Furthermore, the preceding series of summer droughts in 2018, 2019 and 2020 may have left subsoil water reserves not fully restored (Walthert and Meusburger 2025), depriving the trees of a small but notable water source (Pietig et al. under review, Hackmann et al. 2025).

The dendrometer records indicate that radial growth probability declined already at higher REW levels in Douglas fir than in beech. In Douglas fir, this decline occurred roughly in parallel with the decrease in stem water content and sap flow rate, whereas beech sustained growth longer (Figure 3). However, in both species, growth probability started to decline before a REW of ∼0.1, which is when stem rehydration deteriorated, a parameter commonly associated with a turgor-driven decline in stem cambial activity (Zweifel et al. 2016, Peters et al. 2021). Although daily radial growth depends on nighttime rehydration of stem tissues (i.e., turgor), it is also controlled by a complex interplay of carbon source and sink limitations (Mund et al. 2020, Trugman and Anderegg 2024, Gessler and Zweifel 2024) and regulated by hormones (Buttò et al. 2020). Furthermore, dendrometers only capture the phases of growth associated with volumetric expansion (i.e., the turgor-driven cell enlargement), while the carbohydrate-dependent cell wall thickening during cell maturation is not reflected in the measurements (Cartenì et al. 2018, Buttò et al. 2019, Fajstavr et al. 2025). These interacting physiological mechanisms may weaken the direct effect of soil moisture and atmospheric conditions on radial growth.

A key result of our study is the finding that growth started to decline at higher levels of REW and stem rehydration in Douglas fir than in beech (Figures 3 and 4). A consequence of this higher sensitivity is that Douglas fir faced a stronger growth decline in the REW 0.6 – 0.2 range than beech, indicating a considerable drought sensitivity of Douglas fir growth, which seems to be higher than in beech (Leuschner and Meinzer 2024). However, other than beech, Douglas fir resumed growth after drought release in late summer, resulting in both species in a similar number of growth days throughout the entire 2022 vegetation period. While this points at a certain degree of within-season growth resilience, recent long-term growth decline of Douglas fir stands in some regions of the Central European lowlands (Enderle et al. 2024, Cavelier et al. 2025) suggest that the high productivity of this species might not be maintained under climate change. This challenges the widespread belief that Douglas fir is a drought-resistant tree species (Eilman and Rigling 2012, Chakraborty et al. 2019).

### Seasonal development of constraints on stem water relations and growth activity

Despite both species exhibiting declines in stem water content and stem rehydration at similar REW, Douglas fir approached these stem water status thresholds faster in the pure stand than beech (Table 4), which supports our second hypothesis. This can be attributed to Douglas fir exhausting soil water reserves more rapidly due to its higher stand basal area and leaf area index (Paligi et al. 2025), even though the species exhibits an isohydric behavior, which is commonly associated with a more conservative water use (Tardieu and Simonneou 1998, Klein 2014). Furthermore, the conifer showed a similar mid-summer drop in midday leaf water potential (Fig S5) and terminated radial growth earlier than beech, suggesting that Douglas fir did profit only little from its more isohydric strategy in this severe summer, compared to anisohydric beech. This contradicts our third hypothesis. We conclude that – rather than isohydricity – stand structural properties, notably a large stand basal area and high leaf area index, together with the evergreen leaf habit, which enables an early onset of transpiration in spring, are the primary determinants of the water consumption and soil moisture depletion patterns in the Douglas fir stand (Paligi et al. 2025). This confirms that plant morphology (leaf area) and phenology (evergreen habit) can be as important for drought vulnerability as is physiology (Volaire 2018, Kannenberg et al. 2021).

### Reduced drought exposure in the mixed stand

The mixed beech-Douglas fir stand showed a slower soil moisture depletion than either of the pure stands, and the time elapsed to reach REW 0.5, 0.7 or 0.9 was longer, supporting our fourth hypothesis. Both differences in tree water use and in canopy interception may have contributed to the higher soil water availability maintained in the mixed stand. According to an isotopic tracer study in the same stands during our study period, Douglas fir took up water at somewhat greater depth in the pure stand than in the mixed stand (50% water uptake until 40 cm and 28 cm depth, respectively; Hackmann et al. 2025), which may point at more intense root competition in the pure Douglas fir stand compared to the mixture. Beech, in contrast, showed only slightly deeper uptake depths in the mixed compared to the pure stand (34 vs. 31 cm, respectively). These results suggest that Douglas fir in monoculture had to exploit the soil water reserves to greater depth due its overall higher water consumption in comparison to the mixture with beech (Paligi et al. 2025), which had a 12% lower annual water consumption during the dry year and thus likely faced a reduced root competition intensity. It appears that Douglas fir has profited from the admixture of beech, as it showed in mixture a lower reduction in sap flow under drought than in the pure stand (−28% vs. −38%), and its TWD was (up to a certain threshold) lower than in the pure stand (Hackmann et al. 2024, Paligi et al. 2025). Additionally, beech sap flow under drought was unaffected by the admixing of Douglas fir (Paligi et al. 2025), but stem rehydration was improved under drought (Hackmann et al. 2024). This suggests that both species may in a certain way have profited from the mixing. A possible explanation of this pattern is that Douglas fir with higher water consumption per basal area (Paligi et al. 2025) profited from the less water-consuming beech as a neighbor, while beech benefited from partial crown shading by the markedly taller Douglas fir trees in the mixture through reduced transpiration, even though the higher water consumption of the conifer might represent a disadvantage for beech. These findings support the assumption that certain tree species mixtures can stabilize forest stands in drought periods (Forrester 2014, Ammer 2019).

### Limitations of an isohydric behavior as a drought avoidance mechanism

Despite the isohydric behavior of Douglas fir, reflected in a stronger water consumption reduction under drought than in anisohydric beech, the species reached stem water status thresholds (50 to 90% reduction in stem water content and stem rehydration) at similar REW levels as did beech. Moreover, reductions in growth rate were approached earlier despite the more stringent reduction of transpiration. Even more relevant may be the shorter time Douglas fir needs to reach these thresholds. This suggests that being an isohydric, water-demanding species does not represent an overall advantage during drought, as the early depletion of soil moisture may increase the vulnerability to prolonged drought stress.

Our data further challenge the assumption that an isohydric species such as Douglas fir uses soil moisture principally more conservatively to minimize leaf water potential fluctuations (Klein 2014, Tardieu and Simonneau 1998). Contrary to this expectation, Douglas fir exhibited lower (i.e., more negative) midday leaf water potentials than beech during the soil desiccation period (Figure S6), consistent with earlier field observations (Schumann et al. 2024). Further, despite its isohydric regulation, Douglas fir consumed more water than beech even under drought conditions due to its large leaf area (Paligi et al. 2025). Consequently, an isohydric behavior does not necessarily imply the maintenance of a more favorable water status during drought, compared to anisohydric species (Garcia-Forner et al. 2017, Yi et al. 2019). These findings underscore the complexity of species-level drought adaptation and the limitations of isohydricity as a predictive framework for assessing the drought response of trees (Hochberg et al. 2018, Volaire 2018, Paligi et al. 2024).

### Shortcomings of the study and future research directions

In our study, we observed sap flow to continue even at very low REW levels (< 0.05) in both Douglas fir (9% and 14% of the observed maximum in the pure and mixed stand, respectively) and beech (23% and 38% in the pure and mixed stand, respectively). However, since our REW calculation was restricted to the 0-100 cm layer of the mineral soil, it is possible that a significant fraction of root water uptake in such dry periods occurred in the subsoil <100 cm (ca. 9%; Hackmann et al. 2025). Furthermore, the organic layer can considerably contribute to root water uptake; its contribution was estimated at ∼7% of total water uptake in our stands (Hackmann et al. 2025). Future research should extend discrete measurements of midday water potential to continuous potential recordings at foliage, branch and stem levels. This would enable a more comprehensive understanding of how dynamic change in water potential influences the key physiological processes stem water content regulation, sap flow, stem rehydration, and radial growth, as it has been explored in recent studies (Magh et al. 2025, Fajstavr et al. 2025, Peters et al. 2025b).

Since we included only one plot per species combination, our results may reflect site-specific effects. Given that site and stand characteristics can strongly influence drought responses, future studies should extend this research across different species combinations and environmental conditions to assess the broader applicability of our findings (Cai et al. 2024, Niessner et al. 2024, Wankmüller et al. 2024). Furthermore, while the study focused on immediate drought responses, an important next step is to investigate the long-term recovery potential of these species and their capacity to endure prolonged stress periods (Hesse et al. 2023). Understanding species resilience beyond initial drought impacts will be crucial for developing effective forest management strategies in the face of climate change.

### Conclusions

This study examined the co-variation of two measures of stem water status (stem water content and stem rehydration) with sap flow and radial growth dynamics during an extended summer drought for an isohydric conifer (Douglas fir) and an anisohydric broad-leafed species (beech). While both species reached reduction thresholds of SWC and stem rehydration at similar REW, Douglas fir approached these thresholds approximately 200–500 kPa·hours earlier than beech, reflecting the faster soil moisture depletion driven by the species’ higher transpiration rate. Despite its isohydric behavior, Douglas fir exhibited a higher drought-sensitivity of radial growth than beech. These findings confirm that a species’ drought resistance and/or water use strategy is not sufficiently characterized by its degree of isohydricity. A practical consequence for forestry is that fast-growing and water-demanding species may well exacerbate climate change-induced soil water deficits at the stand-level, highlighting the trade-off between maximizing productivity and conserving soil water resources. Mixed-species forests may partly mitigate this negative effect.

Another important finding is that the capacity for nocturnal stem rehydration (which also covers cambium and phloem) was sustained during dry-down longer than stem xylem water content and radial growth. This is important information to be included in the discussion about the factors that limit stem cambial activity during drought. By measuring sap flow, stem water content and diurnal stem diameter variation synchronously, we were able to disentangle key processes underlying the drought response of trees. Such a setting may offer a faster, more precise way to predict tree performance under soil desiccation than water status measurements in the canopy, while being easy to install, of low maintenance effort, and capable of delivering high-resolution data on stem water and growth dynamics. Recognizing more general patterns of co-variation in stem water status, water consumption and growth will require studying a greater diversity of tree species and tree functional types.

## Acknowledgements

SP, CH, JS, MA, CA, DS, and CL acknowledge financial support received from the German Research Foundation (DFG) through the Research Training Group 2300 (316045089/GRK2300). We thank Serena Müller for her outstanding support in coordination and organizing the fieldwork and logistics, which was crucial for a smooth execution of this study. We extend our gratitude to the student helper Lena Zander and to the DAAD RISE-funded interns Jacob Blais, Hailey Stoltenberg and Rachel Einecker for their assistance in the field, and to the fellow tree climbers Klara Mrak, Benjamin Wildermuth and Björn Lüdeke for their help in accessing branches.

## Author contributions

*SP* – study design, conceptualization, collection and processing of sap flow data, data analysis, writing of original draft; *CH* – study design, conceptualization, collection and processing of dendrometer data, data analysis, writing of original draft, *JS* – data analysis; *MA* – conceptualization*; HC* – technical support and supervision in maintenance of sap flow and soil moisture station; *MM* – supervision; *CA* – study design, supervision; CL – study design, supervision, writing of original draft; all authors commented on and approved the final version.

## Data and Materials Availability

The data that support the findings of this study will be provided upon a reasonable request to the first authors SP and CH.

## Supplementary information

### Modeling standardized time to soil desiccation

The data was filtered to capture the period from the peak REW (REW>0.95) in early February to the lowest REW value reached in mid-August (REW<0.05) in all three plots. To investigate the temporal patterns of soil desiccation, we fitted a Shape-Constrained Additive Model (SCAM) with a monotonic penalized decreasing spline to model the relationship between REW and cumulative VPD hours in the timer interval, as implemented in the R package `scam v1.2-12` (Pya 2021). To accommodate plot-specific desiccation patterns, separate smooth functions were used for the three plots. The flexibility and smoothness of the model was balanced by specifying the model with basis dimension 12. This approach allowed us to express the progressive decline of REW over the season and identify the points in time, when specific thresholds were crossed. To analyze temporal desiccation patterns, we extracted the (standardized) time (in kPa hours) elapsed until REW levels of 0.9, 0.7, 0.5, 0.3 or 0.1 were reached in the three plots. These time spans were determined through linear interpolation between the predicted REW levels and the standardized desiccation time.

### Modeling tree water deficit responses to soil desiccation

For statistical analysis, TWD_min_ was normalized by expressing the values of each species as a fraction of 1 (range 0 to 1). For covering the main growing season, the data was restricted to the period from 5^th^ of May to mid-August (in Douglas fir to the end of August).

We investigated the relationship between TWD_min_ and REW by using a generalized additive model (GAM) employing the Tweedie regression family and a logit link function. The tweedie distribution allowed us to account for the many 0 observations that naturally occur in TWD_min_ data. The model included smoothing terms for REW for each neighborhood to capture non-linear relationships with TWD_min_, while also accounting for plot and species-specific differences. Additionally, a random intercept smooth was included to account for unobserved variation among individual trees.

### Sensitivity of sap flow to soil desiccation

The cumulative daily sap flow was calculated, and the data was normalized to enable comparison among trees and species. The data for sap flow was filtered to include data points for the range between saturating soil moisture and maximum soil desiccation. In all plots, maximum soil moisture values exceeded a relative extractable water (REW) value of 0.6. For beech, data before leaf-out (late April) were excluded. Further, the data filtering included only days with mean VPD > 0.5 kPa.

The relationship between sap flow and REW was examined using a generalized additive model (GAM) with a Tweedie regression family and a logit link function. The model included a tensor product smooth for REW (cubic regression spline) and neighborhood (random effect), to capture non-linear relationships with sap flow. This allowed for the modeling of complex relationships between REW and sap flow, while also accounting for differences between plots. Additionally, a random intercept was included to account for variation in sap flow among individual trees.

### Stem water content

The data was filtered to include the period from sensor installation until the end of drought. For Douglas fir in the pure plot, the data started from the 3^rd^ week of March, while for the mixed plot and the pure beech plot, the data started in the 3^rd^ week of April. The data was included until mid-August for beech and end of July for Douglas fir (see Supplementary materials for more details on data filtering). The chosen period was characterized by higher relative extractable water across all plots with a minimum value of 0.62 in the pure Douglas fir plot. After data filtering, the stem water content values were normalized for each tree by scaling them between their maximum (1) and minimum values (0) to facilitate comparison between trees and plots.

We examined the relationship between stem water content (SWC) and relative extractable water (REW) using a generalized additive model (GAM) under the framework of the tweedie regression family with a logit link function. The model was specified with a cubic regression smoothing splines for REW, separately for each neighborhood, with five knots to capture non-linear relationships with SWC, while also accounting for plot-specific differences. A random intercept was included to account for unobserved variation in SWC among individual trees. Since the measurements were taken at two different positions (inner and outer) with different results, another random intercept for measurement position nested in tree was included.

#### Data processing and filtering criteria

To ensure robustness and ecological relevance of our analysis, we applied targeted data filtering criteria tailored to each physiological response variable, thus reflecting realistic drought responses under natural conditions. In all cases, the earliest data period to be included in the analysis differed by variable and plot type, starting on 21^st^ March for stem water content in pure Douglas fir and on May 10^th^ for growth probability, sap flow and TWD in beech according to the leaf-out phenology. The latest data points extended until the major precipitation event on August 13^th^.

We employed variable specific data filtering procedures to reflect the unique responses of each variable to phenological or climatic conditions. For sap flow, we excluded days with a mean vapor pressure deficit (VPD) below 0.3 kPa. This threshold was chosen because sap flow is strongly influenced by VPD, and at very low VPD, tree water use is minimal. For the tree water deficit (TWD) of Douglas fir, data selection was limited to the period up to the end of July. This decision was based on the likelihood of TWD recovery in the species after the small rain events at the end of July. These events potentially led to stem rehydration, thus explaining the lower TWD observed. This was only the case in Douglas fir and not in beech, which is plausible, as Douglas fir as an isohydric species exhibits stricter stomatal closure under drought conditions, apparently prioritizing stem rehydration over water use through stomatal opening (Hammond 2020, Bi et al. 2022).

Nevertheless, across all species, plot types, and variables, our dataset predominantly covered relative extractable water (REW) values > 0.6, ensuring a broad representation of soil moisture conditions. The only exception was the pure Douglas fir plot for TWD, where the dataset included REW values starting from 0.41. However, this does not compromise the analysis, as these periods still included sufficient intervals with zero TWD, ensuring a reliable representation of drought-induced water deficit dynamics. By implementing these filtering criteria, we optimized the dataset to reflect biologically meaningful thresholds, while maintaining comparability across species and stand compositions.

#### Supplementary figures

**Figure S1.**
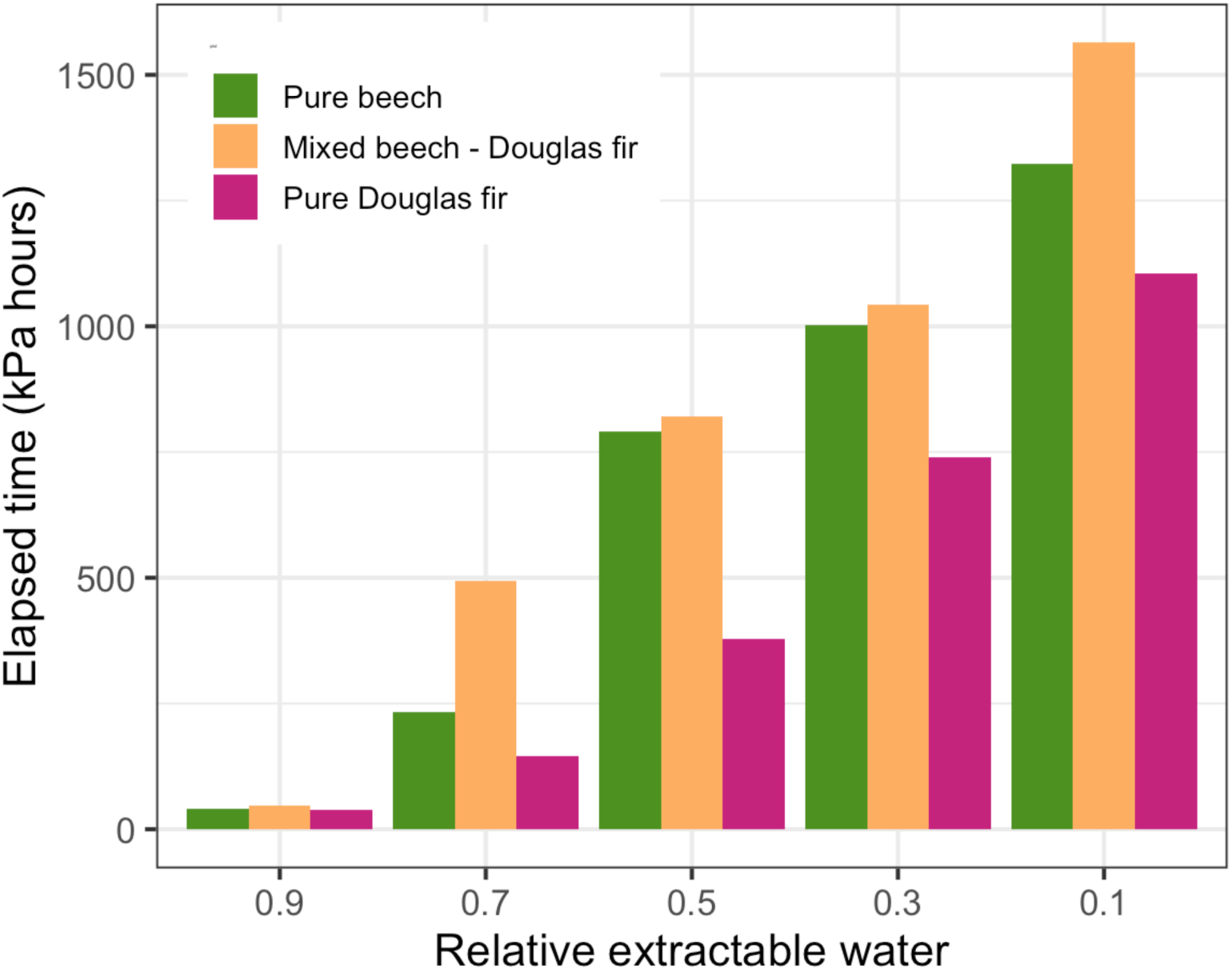
Time elapsed in the pure beech (green), pure Douglas fir (violet) and the mixed beech-Douglas fir stand (orange) to reduce REW to 0.9, 0.7, 0.5, 0.3 or 0.1 during the summer drought in 2022.

**Figure S2.**
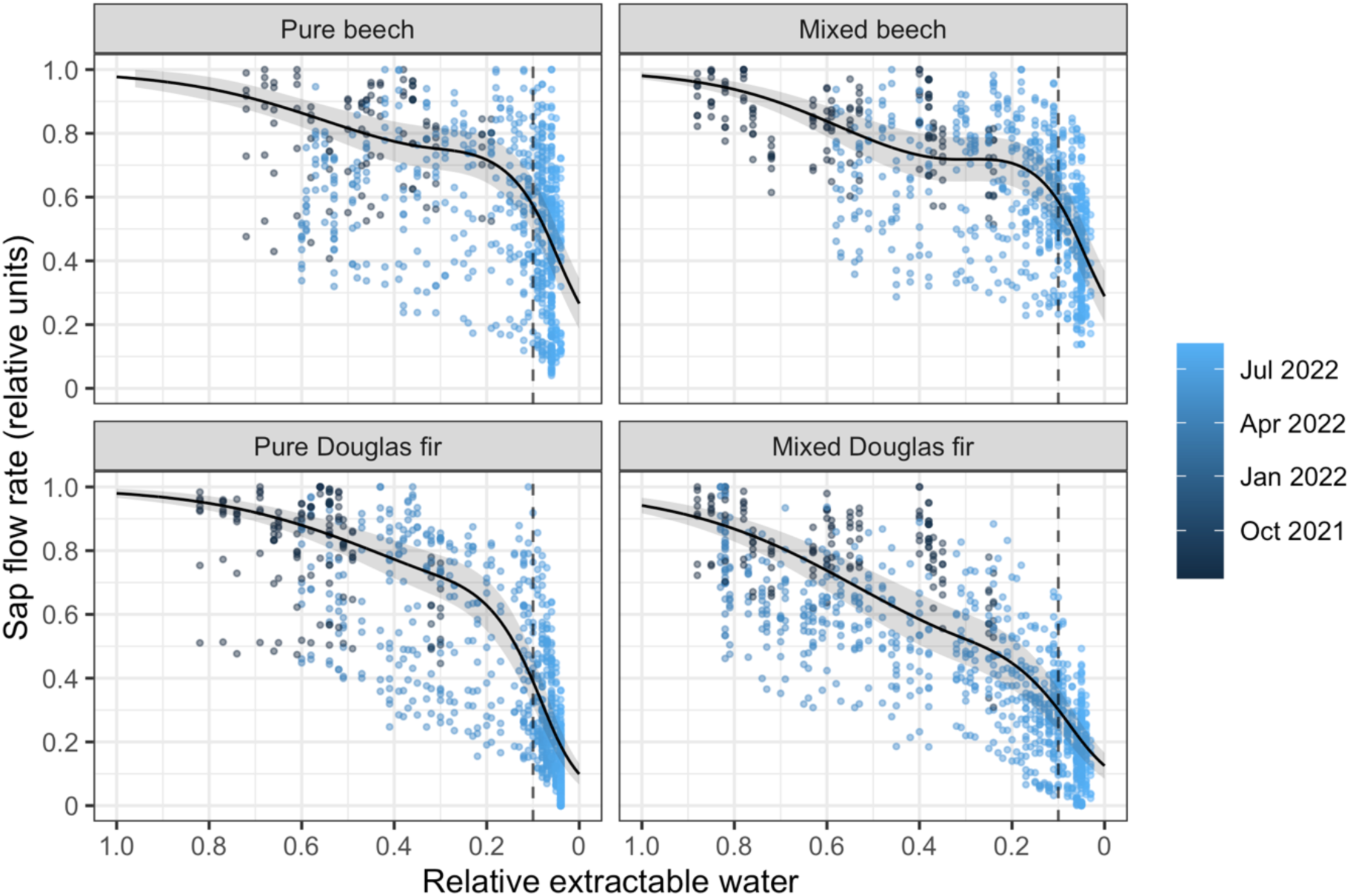
Daily totals of sap flow (given in relative units) in their dependence on relative extractable water (REW) in the pure beech, pure Douglas fir and mixed beech-Douglas fir stand during the soil dry-down in summer 2022 (April – August) and additional data from the drying phase in 2021 (July 10th – September 10th). The color of the dots gives the month of measurement. Each data point represents a daily total obtained by averaging over eight trees per species in each stand. The solid black curves depict Generalized Additive Model (GAM) fits as a tensor product of REW and plot type. The shaded region indicates the 95% confidence interval. The vertical dashed lines denote REW 0.1.

**Figure S3.**
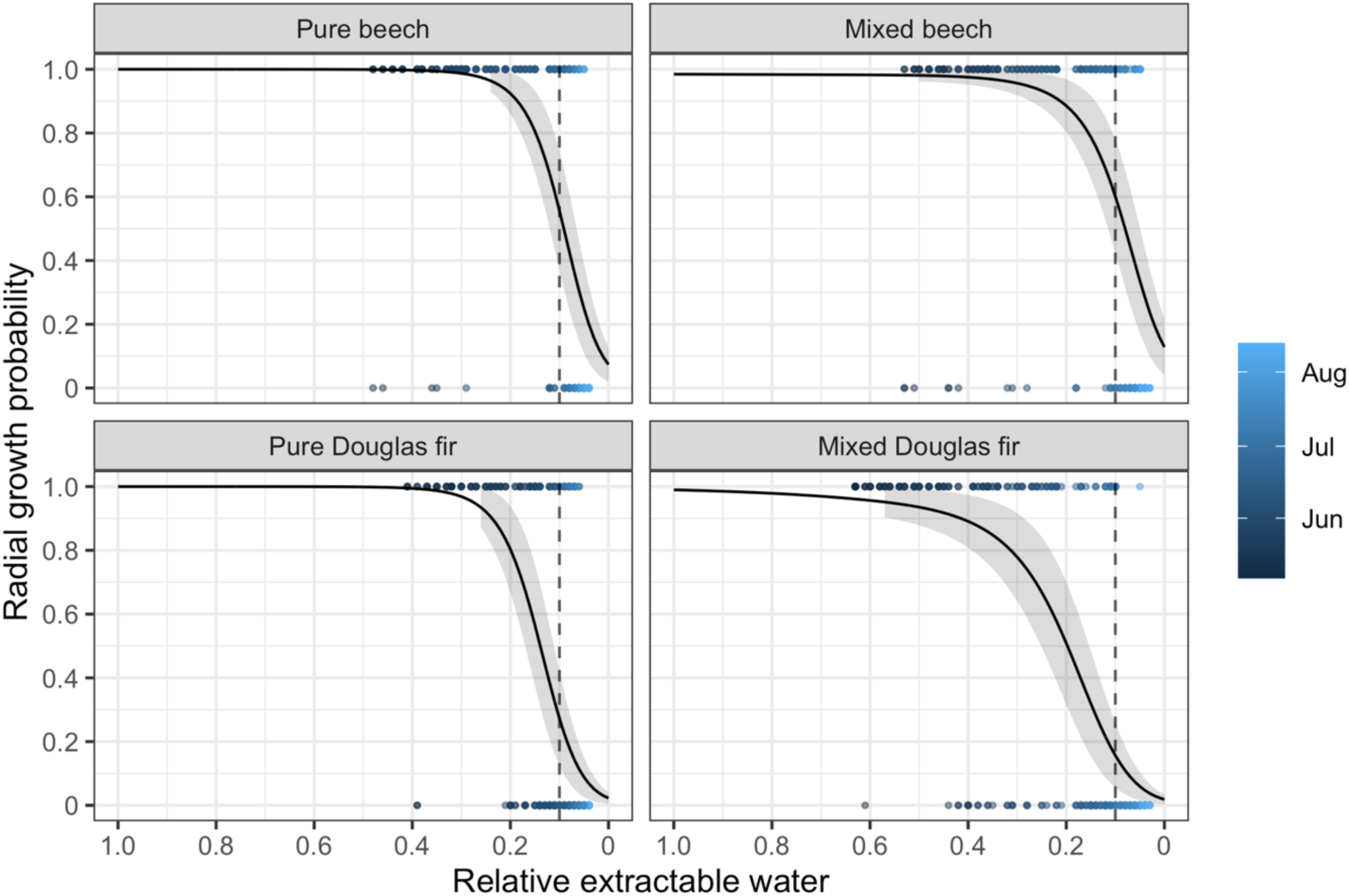
Daily radial growth (given as growth probability according to a GAM, either 1 or 0) in its dependence on relative extractable water (REW) in the pure beech, pure Douglas fir and mixed beech-Douglas fir stand during the soil dry-down in summer 2022 (April – August). The color of the dots gives the month of measurement. Each data point represents a daily growth probability as obtained by averaging over four trees per species in each stand. The solid black curves depict Generalized Additive Model (GAM) fits as a tensor product of REW and plot type. The shaded region indicates the 95% confidence interval. The vertical dashed lines denote REW 0.1

**Figure S4.**
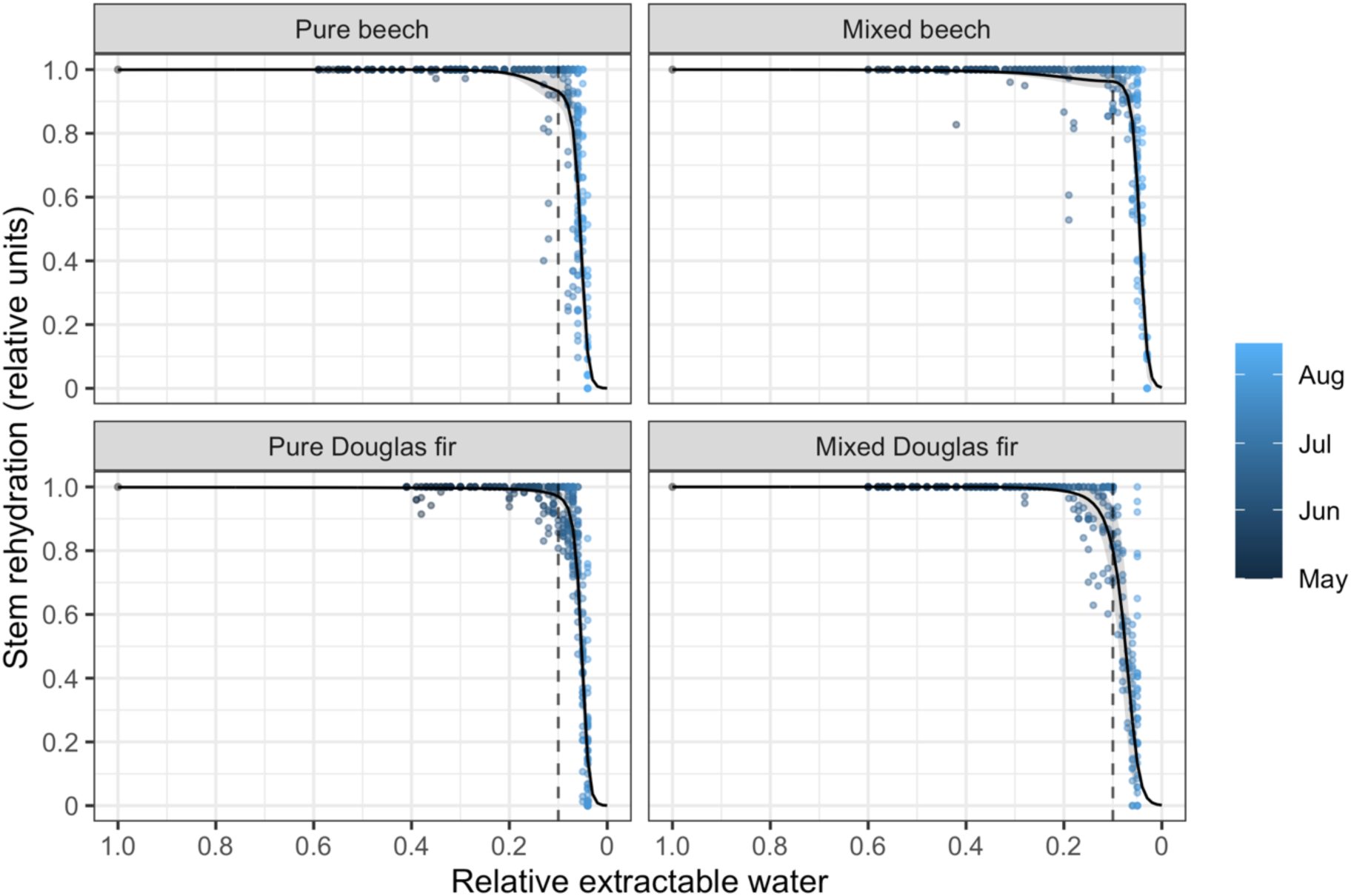
Daily minima of stem rehydration (given in relative units) in their dependence on relative extractable water (REW) in the pure beech, pure Douglas fir and mixed beech-Douglas fir stand during the soil dry-down in summer 2022 (April – August). The color of the dots indicates the month of measurement. Each data point represents a daily maximum stem rehydration obtained by averaging over four trees per species in each stand. The solid black curves depict Generalized Additive Model (GAM) fits as a tensor product of REW and plot type. The shaded region indicates the 95% confidence interval. The vertical dashed lines denote REW 0.1.

**Figure S5.**
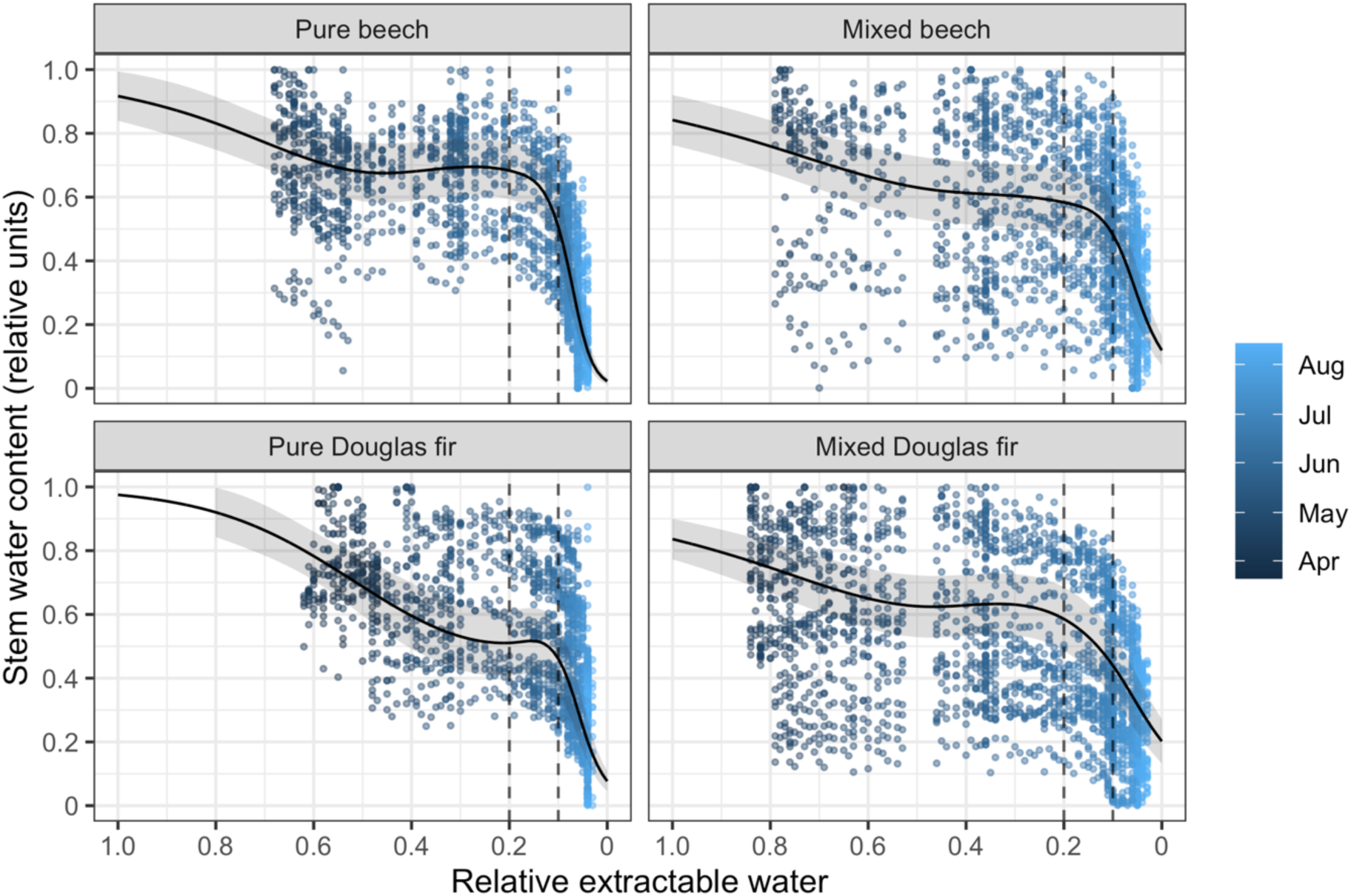
Daily maxima of stem water content (SWC, given in relative units) in their dependence on relative extractable water (REW) in the pure beech, pure Douglas fir and mixed beech-Douglas fir stand during the soil dry-down in summer 2022 (April – August). The color of the dots gives the month of measurement. Each data point represents a daily maximum SWC as calculated by averaging over the outer and the inner position in the bark (0.5 and 1.5 cm of sapwood; n = 8 trees per species in each stand). The solid black curves depict Generalized Additive Model (GAM) fits as a tensor product of REW and plot type. The shaded region indicates the 95% confidence interval. The vertical dashed lines denote REW 0.1 and REW 0.2.

**Figure S6.**
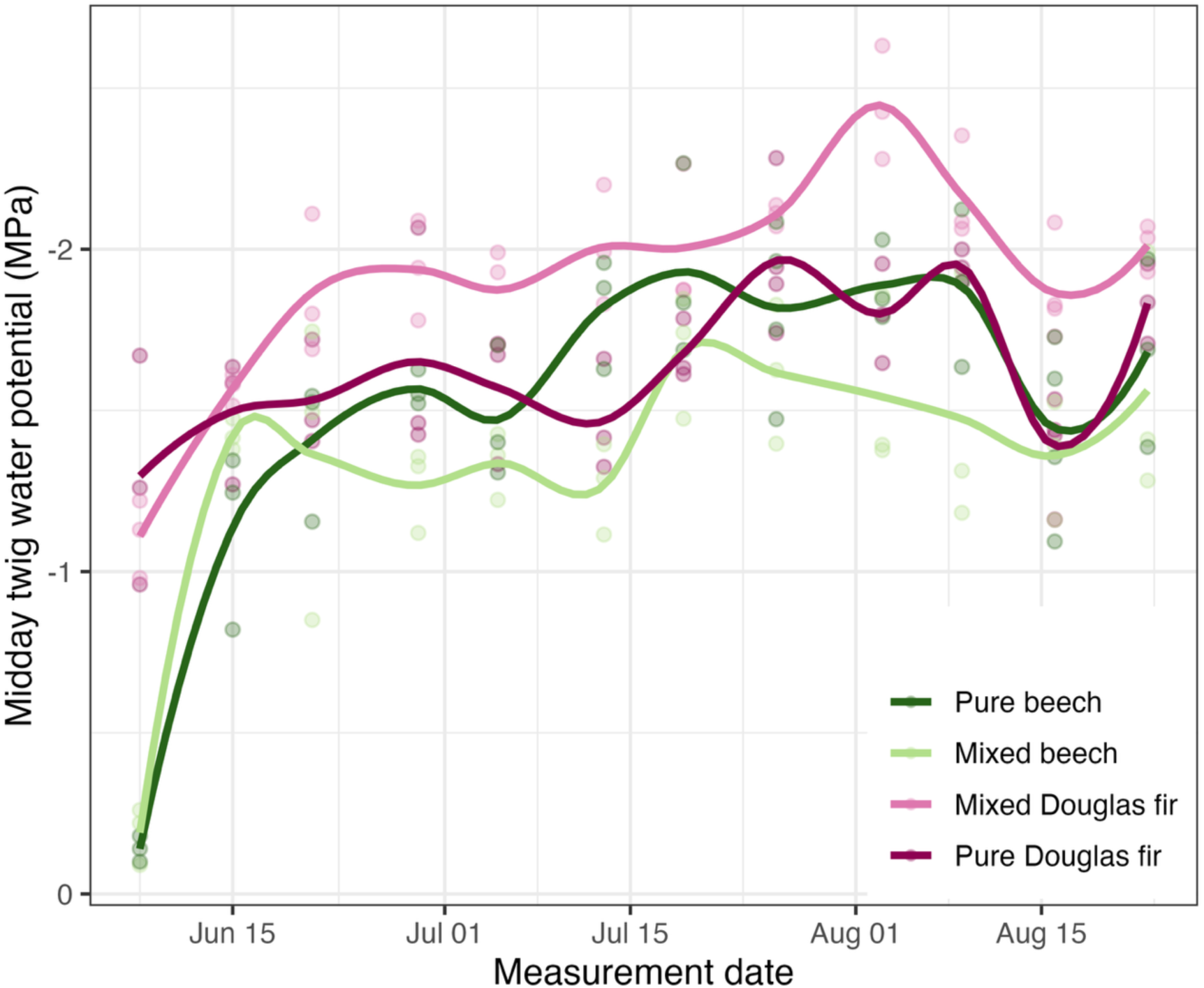
Seasonal variation in midday twig water potential (MPa) across different neighborhood types: Pure beech (light green), mixed beech (dark green), pure Douglas Fir (light pink) and mixed Douglas fir (dark pink). Solid lines represent locally smoothed trends using a loess fit, while individual points correspond to measured values.

